# Inhibition of gonadotropin-stimulated steroidogenesis by the erk cascade

**DOI:** 10.1101/853648

**Authors:** Rony Seger, Tamar Hanoch, Revital Rosenberg, Ada Dantes, Wolfgang E. Merz, Jerome F. Strauss, Abraham Amsterdam

## Abstract

LH and FSH are two important hormones in the regulation of granulosa cells. Their effects are mediated mainly by cAMP/PKA signaling, bit the activity of the extracellular signal-regulated kinase (ERK) signaling cascade is elevated as well. We studied the involvement of the ERK cascade in LH and FSH-induced steroidogenesis in two granulosa-derived cell lines, rLHR-4 and rFSHR-17, respectively. We found that stimulation of these cells with the appropriate gonadotropin induced ERK activation as well as progesterone production, downstream of PKA. Inhibition of ERK activity enhanced gonadotropin-stimulated progesterone production, which was correlated with increased-expression of the steroidogenic acute regulatory (StAR) protein, a key regulator of progesterone synthesis. Therefore, it is likely that gonadotropin-stimulated progesterone formation is regulated by a pathway that includes PKA and StAR, and this process is downregulated by ERK, due to attenuation of StAR expression. Our results suggest that activation of PKA signaling by gonadotropins not only induces steroidogenesis, but also activates downregulation machinery involving the ERK cascade. The activation of ERK by gonadotropins as well as by other agents, may be a key mechanism for the modulation of gonadotropin-induced steroidogenesis.

## Introduction

Gonadotropic hormones, follicle stimulating hormone (FSH) and luteinizing hormone (LH), play a crucial role in controlling reproductive function in males and females. These hormones are released from the pituitary and induce pleotropic effects in various cells of the reproductive system including ovarian granulosa cells (LH and FSH), theca interna cells (LH), testicular Sertoli cells (FSH), and Leydig cells (LH) (1-3). In the ovary, the main effects of both LH and FSH are stimulation of estradiol and progesterone, which in turn, control the reproductive cycle (reviewed in (4)). In the past years, progesterone production in ovarian granulosa cells have been studied in details. Among others, it was shown that the gonadotropins exert their stimulatory activity via the G-protein coupled receptors (GpCR) either LH-receptor and FSH-receptor, to which each gonadotropin bind specifically. Upon binding, both receptors stimulate the Gs protein, which then induces elevation of intracellular cAMP activates via the membrane-associated adenylyl cyclase, causing an (5). In turn, the cAMP serves as a second messenger for the upregulation of the steroidogenic acute regulatory protein (StAR) and the cytochrome P450 (P450scc) enzyme system (reviewed in (6,7)).

It is clear that the cAMP-StAR axis is clearly the main pathway that regulate steroidogenesis. However, several other signaling processes have been implicated in this regulation as well. Thus, it was shown that steroidogenesis is regulated by desensitization of the gonadotropin receptor (3). In addition, gonadotropin receptors can activate several other signaling processes, including calcium mobilization, activation of the phosphoinositol pathway, and stimulation of chloride ion influx (reviewed in (8)). In addition, G-protein coupled receptor kinase phosphorylation of the gonadotropin receptors, the adaptor protein arrestin and massive internalization of the receptors are thought to play a role in the downregulation of gonadotropin signaling. However, the link of these pathways to steroidogenesis is not well-proven yet (5). Nonetheless, since all these processes are important for the regulation of the receptors and their downstream activities (9), it is likely that additional mechanisms participate in the rapid attenuation of gonadotropin signals and in the regulation of steroidogenesis.

The Extracellular-signal regulated kinases (ERKs) is a group of signal transduction protein kinases composed of three members (p42ERK2, p44ERK1, p46ERK1b). This group belongs to the family of the signaling mitogen-activated protein kinases (MAPKs). Upon extracellular stimulation, the ERKs are activated by a network of interacting proteins that direct the signals into a multi-tier protein kinase cascade, mainly Raf MEK ERK and MAPKAPKs (reviewed in (10,11)). The activated ERKs in turn can phosphorylate and activate target regulatory proteins in the cytoplasm (e.g. PLA2) and in the nucleus (e.g. Elk1). These effectors then govern various cellular processes, mainly proliferation and differentiation. It is presently known that these kinases also participate in other processes including the control of cellular morphology carcinogenesis and more (11).

In previous studies it has been shown that ERK is significantly activated (2-5 fold) upon stimulation of ovarian granulosa cell with LH and FSH (12,13). This activation is dependent on cAMP and PKA, as it is mimicked by elevation of intracellular cAMP, and attenuated by inhibitors of PKA. In the present work, we found that both gonadotropins, LH and FSH induce ERK activation and progesterone production. We also showed that in immortalized granulosa cell lines, these effects are mediated by cAMP. It is important to mention, that these cell lines consist of homogeneous populations with respect to the LH receptor, unlike the follicular granulosa cells which were previously used to show the effect (14). Interestingly, inhibition of ERK using a MEK inhibitor caused an elevated gonadotropin-cAMP-induced progesterone production, while activation of ERK inhibits this process. Moreover, we found that MEK inhibitor elevated the intracellular content of StAR, which operates downstream of cAMP. Therefore, it is possible that the inhibitory effect of the ERK on steroidogenesis may be mediated by reduced StAR expression. Thus, we concluded that gonadotropin-induced progesterone formation is regulated by PKA. The latter induces not only StAR expression, but also a counteracting down-regulating mechanism. These two mechanisms are acting simultaneously due to regulation of the activated ERK, which reduces StAR expression.

## Experimental Procedures

### Stimulants, inhibitors antibodies and other reagents

Human FSH (hFSH) human LH (hLH) and human chorionic gonadotropin (hCG) were kindly provided by the NIH and Dr. Parlow. Deglycosilated hCG was enzymatically prepared as previously described (15). Mouse monoclonal anti-diphospho ERK (anti-active ERK/MAPK) antibodies (DP-ERK Ab), and anti-general ERK antibody were obtained from Sigma, Israel (Rehovot, Israel). Anti-C-terminal ERK1 antibody (C16) was purchased from Santa Cruz. Polyclonal antibodies to human StAR were raised in rabbit (16). Alkaline phosphatase, horseradish peroxidase and flourescein conjugated secondary antibodies were purchased from Jackson ImmunoResearch laboratories Inc. (West Grove, Pennsylvania). PD98059 and U0126 were purchased from Calbiochem (San Diego). H89, Forskolin, 8-Br-cAMP were obtained from Sigma (St. Louis).

### Cell lines

rLHR-4 cell line was established by cotransfection of rat preovulatory granulosa with mutated p53 (Val135-p53) Ha-*ras* genes and plasmid expressing the rat LH/CG-receptor (17). The rFSHR-17 cell line was established by immortalization of preovulatory rat granulosa cells via cotransfection of primary cells with SV40 DNA and an HA-*ras* gene. Cells were transfected with plasmid expressing the rat FSH receptor (18). The cells were maintained in F12/DMEM medium (1:1) containing 5% fetal calf serum.

### Stimulation and harvesting of cells

Subconfluent cultures were serum-starved for 16 h, and subsequently incubated for selected time intervals with the indicated agents in the presence or absence of various inhibitors. Following stimulation, cells were washed twice with ice-cold phosphate buffered saline, once with Buffer A (50 mM β-glycerophosphate, pH 7.3, 1.5 mM EGTA, 1 mM EDTA, 1 mM DTT, and 0.1 mM sodium vanadate, (19)), and were subsequently harvested in ice-cold Buffer A + proteinase inhibitors (19). Cell lysates were centrifuged at (20,000xg, 20 min). The supernatant was assayed for protein content and subjected to a Western blot analysis or to immunuprecipitation as below. For the detection of StAR protein, cells were lysed in RIPA Buffer (19) and subjected to Western blot analysis.

### Transfection of PKI and ERK plasmids into cells

The rLHR-4 and rFSHR-17 cells were grown in Dulbecco’s modified Eagle’s medium (DMEM) supplemented with 10% fetal calf serum (FCS) up to 70% confluency. The plasmids used were RSV-PKI and RSV-PKI^mutant^, (20)) (a generous gift from Dr. R.A.Maurer, Oregon Health Sciences University, Portland), and pGFP-ERK2 (21). The plasmids were introduced into the two cell types using Lipofectamine (Gibco-BRL) according to the manufacturer instruction. About 15-20% transfection was observed in the two cell lines using a Zeiss florescent microscope. After transfection, the rLHR-4 cells were grown in DMEM+10% FCS for 6 hours and then starved in DMEM+0.1% fetal calf serum for additional 14 hours. The rFSHR-17 cells were grown in DMEM+10% FCS for 20 hours. The transfected cells were then stimulated and harvested as above.

### Western Blot Analysis

Cell supernatants, which contained cytosolic proteins were collected, and aliquots from each sample (30 *µ*g) were separated on 10% SDS-PAGE followed by Western blotting with the appropriate antibodies. Alternatively, immunoprecipitated proteins were boiled in sample buffer and subjected to SDS-PAGE and Western blotting. The blots were developed with alkaline phosphatase or horseradish peroxidase-conjugated anti-mouse or anti-rabbit Fab antibodies.

### Determination of ERK activity by phosphorylation

Cell supernatants (200 *µ*g proteins) were subjected to immunoprecipitation with monoclonal anti-ERK C-terminal antibodies (C16, Santa Cruz) as described above. During the final step of immunoprecipitation, pellets were washed with buffer A, resuspended in 15 μl of buffer A, and incubated (20 min, 30°C) with 5 μl of 2 mg/ml myelin basic protein (MBP) and 10 μl of 3x reaction mix (30 mM MgCl_2_, 4.5 mM DTT, 75 mM β-glycerophosphate, pH 7.3, 0.15 mM Na_3_VO_4_, 3.75 mM EGTA, 30 *µ*M calmidazolium, 2.5 mg/ml bovine serum albumin and 100 *µ*M [γ^32^P]-ATP (2 cpm/fmol)). The phosphorylation reactions were terminated by addition of sample buffer and boiling (5 min) and the samples were analyzed by SDS-PAGE and autoradiography as previously described (19).

### Progesterone assay

Progesterone secreted into the culture medium was assayed by radioimmunoassay as previously described (22).

### Localization of StAR protein by immunofluorescence

cells were cultured on 24×24 mm cover glasses placed in 35 mm plastic tissue culture dishes. Cells were fixed with 3% paraformaldehyde subsequent to 24 h incubation at 37°C with the appropriate stimulants and visualized in a Zeiss florescent microscope following incubation with 1:200 dilution of antiserum to human StAR and goat anti-rabbit antibodies conjugated to floresceine. For negative controls cells were incubated with non-immune rabbit serum followed by the second antibodies.

## Results

Stimulation of granulosa cells with the gonadotropins LH or FSH induces several cellular processes, including de-novo synthesis of steroid hormones. Here we undertook to study the intracellular signaling leading from the LH and FSH receptors to the gene transcription that regulate progesterone production. For this purpose, we used two distinct granulosa cell lines expressing either LH/CG or FSH receptors: rLHR-4 and rFSHR-17 respectively. It had been previously shown that addition of each of the gonadotropins to their corresponsing cells stimulats cAMP production, activation of PKA and induction of steroidogenesis ((18) and data not shown). Since the ERK cascade was implicated in the signaling of G protein-coupled receptors (23), we first examined whether the ERK cascade is also activated in the rLHR-4 and rFSHR-17 cell lines.

### Activation of ERK by hCG, Deglycosylated hCG (dghCG) and cAMP in rLHR-4 cells

Serum-starved rLHR-4 cells were stimulated with hCG, which signals via the LH receptor (3). The activating phosphorylation of the ERK on its TEY motif was assessed using a Western blotting with DP-ERK Ab (24). Considerable staining of three bands at 42, 44 and 46 kDa (ERK2, ERK1 and ERK1b respectively (19)) was detected in the resting, non-stimulated, cells. The intensity of staining of ERK2 and ERK1 was enhanced (∼5 fold) 5-20 min after the addition of hCG, and remained high (∼3 fold) up to 60 min after stimulation. The appearance of p46 ERK1b is of particular interest in these cells, because although ERK1b has been reported to exist in rat (19), its abundance in human cells is usually very small as compared to ERK1 and ERK2. Interestingly, the basal activity of ERK1b in rLHR-4 cells was as high as that of ERK1, but it was only modestly increased (5-20 min, ∼2 fold), and it declined back to basal level 40 min later. The kinetics of activation, which are different from that of ERK1 and ERK2, suggests a differential mode of ERK1b-regulation as recently demonstrated in EJ cells (19).

We next examined LH, which like hCG, specifically acts via the LH-receptors. The effect of LH on ERK activity was essentially the same as that of hCG under all conditions examined (data not shown). We then used deglycosylated hCG (dghCG) that has previously been reported to maintain the same affinity to the LH-receptor as the intact hormone but retains only a residual activity for stimulation of steroidogenesis (25). When added to the cells, it did cause activation of ERKbut to a much lesser degree than the activation achieved by the intact hormone (2,5 fold activation 20 min after dghCG treatment as compared to 4.5 fold 20 min after hCG treatment (Fig. 1)). Since LH and hCG are known to operate via Gs and cAMP (25), we examined whether ERK activation is cAMP-dependent. Indeed, both 8-Br-cAMP, and forskolin, which activate adenylyl cyclase and as a consequence induce cAMP production, also significantly activated ERK phosphorylation in the rLHR-4 cells (data not shown). These results indicate that the hCG-induced ERK activation is probably dependent on cAMP elevation.

**Fig. 1.**
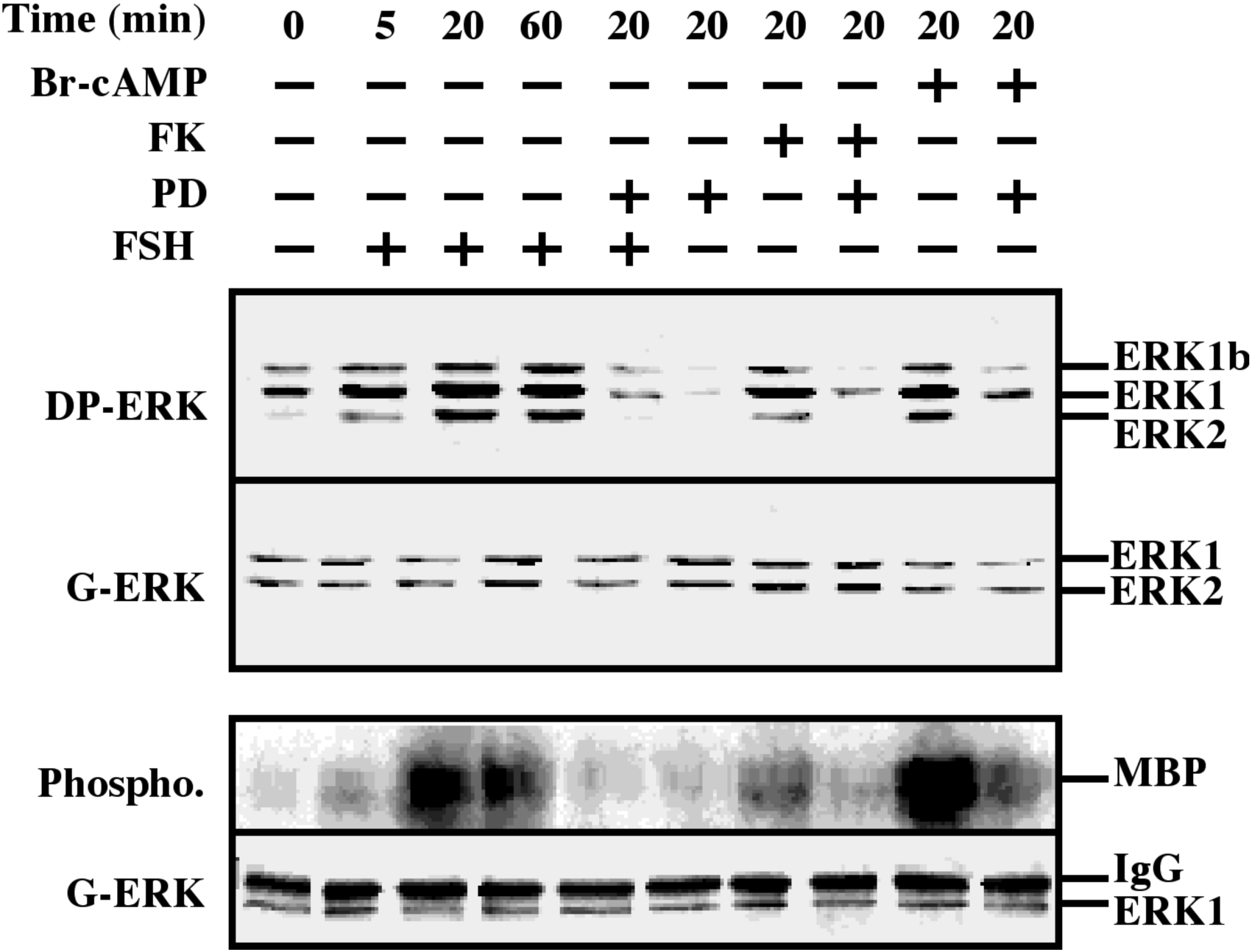
Activation of ERK/MAPK by hCG and dghCG in rLHR-4 cells. rLHR-4 cells were serum-starved for 16 h, and then stimulated with hCG (3 iu/ml) with or without PD 98059 (PD, 15 min prestimulation, 25 *µ*M), with PD98059 (25*µ*M) alone or with dghCG (3 iu/ml) for the indicated times. Cytosolic extracts (50 *µ*g) were subjected to immunoblotting with DP-ERK (upper panel) or with anti-general ERK antibody (G-ERK, second panel). Alternatively, the cytosolic extracts were subjected to immunoprecipitation with anti-C terminal ERK1 antibody (C16) followed by *in vitro* phosphorylation of MBP as described in Material and Methods (3^rd^ panel, Phospho.). The amount of immunoprecipitated ERK for the phosphorylation reaction was determined by Western blotting with the anti-general ERK antibody (bottom panel). The position of ERK2, ERK1 and ERK1b, MBP and IgG is indicated. Each of these experiments was reproduced at least three times.

ERK phosphorylation of the TEY motif is usually correlated to its activation, but mainly reflects MEK activity. In order to assess whether the phosphorylation also correlates with the activity of ERK itself, we resorted to an ERK kinase assay. This was performed by immunoprecipitation of ERK with anti-C-terminal antibody, followed by phosphorylation of the general, non-specific substrate - myelin basic protein (MBP (19)). As expected, we found that the activity of ERK correlated nicely with the regulatory phosphorylation (Fig. 1, bottom two panels), verifying that the 4-5-fold activation of ERK activity in rLHR-4 cells upon gonadotropin stimulation. Tis point was also proven by using the MEK inhibitor, PD98059, which reduced the activity of ERK. This was seen both in hCG-stimulated and non-stimulated activity of ERK to below basal levels. A similar reduction was observed for dghCG-, forskolin- and 8-Br-cAMP-stimulated activity of ERK (Fig. 1 and data not shown). Staining with an anti-general ERK antibody (7884) which recognizes both ERK1 and ERK2 revealed that none of the treatments caused any significant change in the total amount of the ERKs (Fig. 1; G-ERK).

### Activation of ERK by FSH and cAMP in rFSHR-17 cells

We then tested whether FSH is able to stimulate ERK activity as much as LH. This was done using the FSH expressing rat granulosa-derived cell line, rFSHR-17. Indeed, using Western blot, we found that there was considerable detection of all three ERK isoforms, ERK2, ERK1 and ERK1b, in extracts of serum-starved cells. This staining was enhanced upon FSH treatment of the cells, in kinetics that were slightly slower than the kinetics of hCG stimulation in rLHR-4 cells (Fig. 2 upper lanes). The staining of the three ERK isoforms was enhanced 5 min after FSH stimulation, peaked (5-fold above basal level) at 20 min, after stimulation and slightly decreased at 60 min. Importantly, also in these cells, the cAMP stimulating agents, forskolin and 8-Br-cAMP, enhanced the phosphorylation of the three ERK isoforms (3 and 5 fold above basal level, respectively). Again, as expected, the treatments did not cause any change in the amount of the ERK isoforms as judged by general anti-ERK antibody. These results confirm that the changes detected by the DP-ERK Ab are indeed due to changes in ERK phosphorylation and not due to induction of ERK expression. Next, we examined ERK activity using the kinase assay described for the rLHR-4 cells using MBP as a substrate. We indeed found (Fig. 2, bottom) that not only ERK phosphorylation but also ERK activity was stimulated by FSH, forskolin and 8-Br-cAMP similarly were reduced by PD98059. These results show that both LH and FSH receptors can transmit signals to the ERK pathway via cAMP in the examined cell lines.

**Fig. 2.**
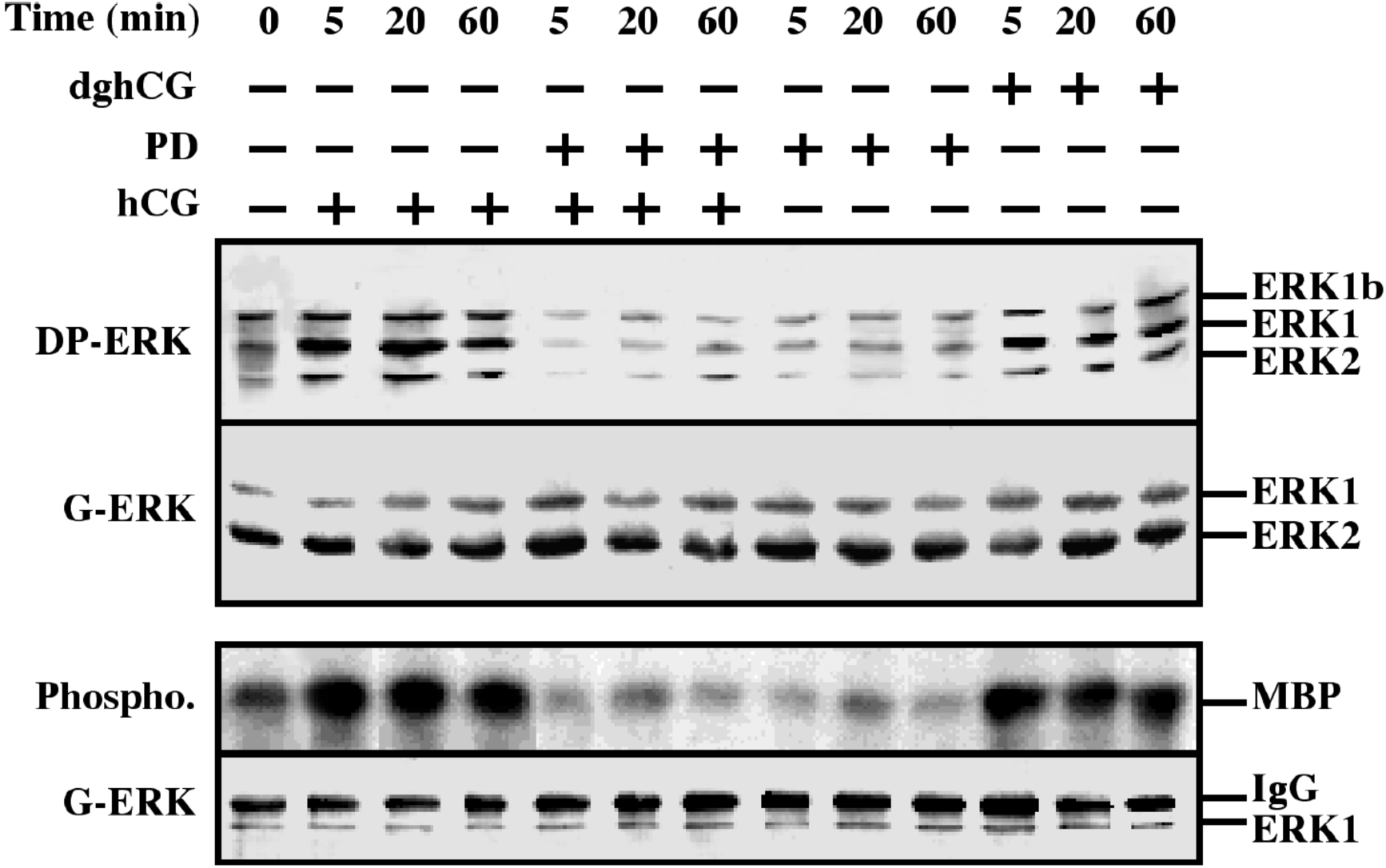
Activation of ERK/MAPK by FSH/cAMP in rFSHR-17 cells. RFSHR-17 cells were serum-starved for 16 hr s and then stimulated with FSH (3 iu/ml) with forskolin (FK, 50 *µ*M), with 8-Br-cAMP (Br-cAMP, 50 *µ*M) with or without PD 98059 (PD, 15 min prestimulation, 25 *µ*M), or with PD 98059 (25 *µ*M) alone for the indicated times. Cytosolic extracts (50 *µ*g) were subjected to immunoblotting with DP-ERK (upper panel) or with anti-general ERK antibody (G-ERK, second panel). Alternatively, the cytosolic extracts were subjected to immunoprecipitation with anti-C terminal ERK1 antibody (C16) followed by *in-vitro* phosphorylation of MBP as described under Material and methods (phospho, 3^rd^ panel). The amount of immunoprecipitated ERK for the phosphorylation reaction was determined by Western blot with the anti-general ERK antibody (bottom panel). The position of ERK2, ERK1 and ERK1b, MBP and IgG is indicated. Each experiment was reproduced at least three times.

### PD98059 stimulates FSH and hCG-induced steroidogenesis

One of the main cellular processes stimulated by gonadotropins in granulosa cells is stereoidogenesis (26). Indeed, we found that progesterone production is significantly increased 24 and 48 h after LH stimulation of rLHR-4 cell line (Fig. 3A). hCG had a similar effect to that of LH (data not shown), while dghCG had a very small effect. Importantly Forskolin caused a two-fold greater induction of progesterone production than LH indicating a role of cAMP in this process. In order to examine whether activated ERK is involved in the induction, we incubated the rLHR-4 cells with the MEK inhibitor, PD98059. We found that the inhibitor had no effect by itself on progesterone production in rLHR-4 cells. However, when the cells were incubated with PD98059 for 15 min prior to LH stimulation there was a 3-fold increase in LH-induced progesterone production (Fig. 3), simultaneously with a complete abolishment of ERK activity (Fig. 1). A Similar effect was observed when the MEK inhibitor was added prior to stimulation of the cells with forskolin (Fig. 3), hCG, and 8-Br-cAMP (data not shown). This effect was similar also for FSH, as similar to the rLHR-4 cells, the MEK inhibitor increased steroidogenesis in rFSHR-17 cells. Thus, in these cells, FSH and forskolin caused a significant elevation of progesterone production after 24 and 48 h, which was dramatically amplified by the addition of PD98059. To further prove this point, we used ERK activation rather than ERK inhibition. In contrast to the induction by MEK inhibitor, TPA, which is a known activator of the ERK (27) had a negative effect on the forskolin-induced production of progesterone in both cell lines after 24 and 48 hours. Taken together, these results indicate that the ERK signaling cascade has inhibitory effect on gonadotropin and cAMP-stimulated progesterone production.

**Fig. 3.**
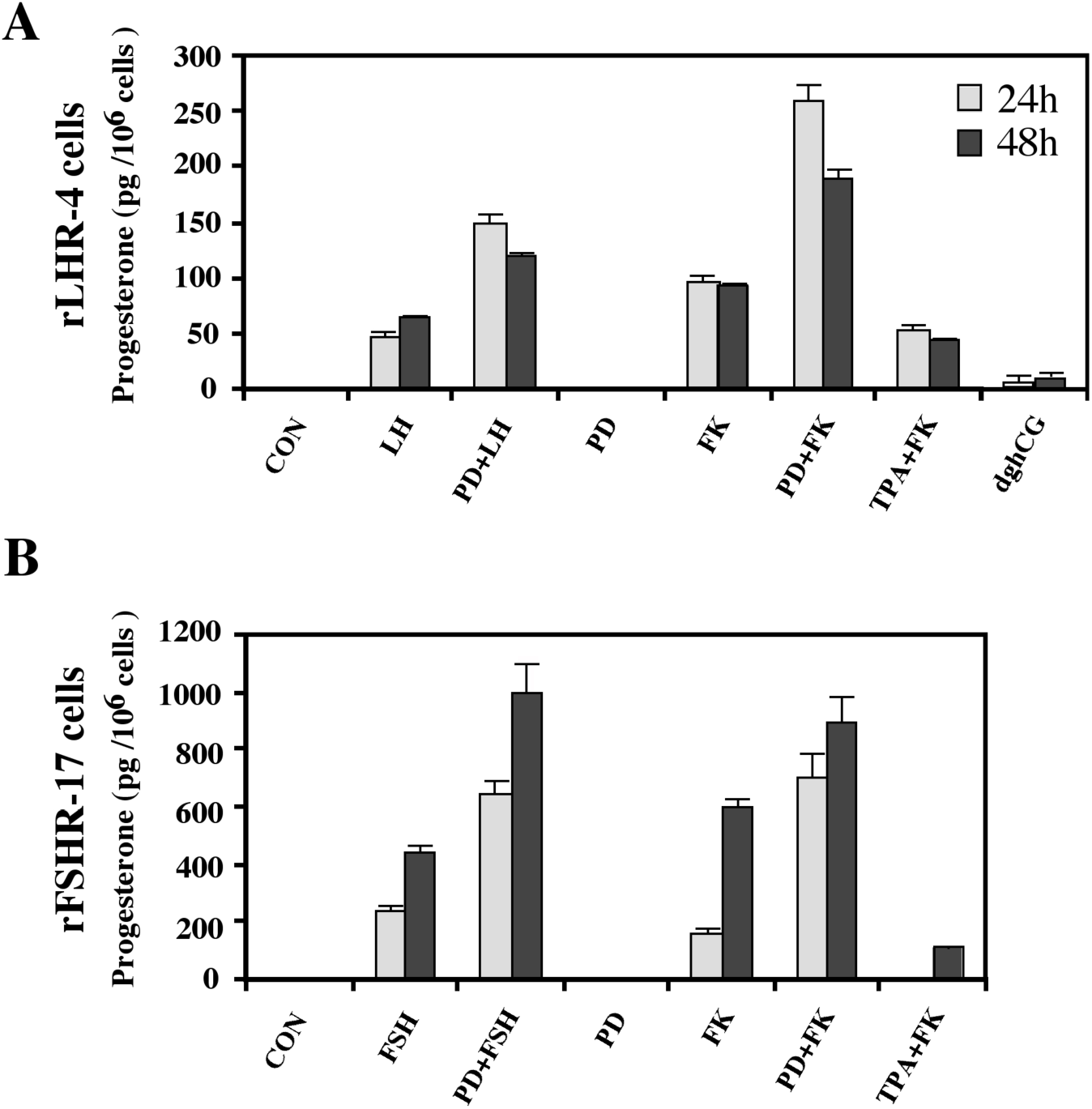
Enhancement of progesterone production by MEK inhibitor, gonadotropins and cAMP-stimulated rLHR-4 and rFSHR-17 cells. Subconfluent cultures were treated with PD98059 alone (PD, 25 *µ*M), hCG (3 iu/ml), hFSH (3iu/ml) dghCG (3 iu/ml), forskolin (FK, 50 *µ*M) TPA (100 nM) or the same reagent with PD98059 as indicated for 24 h or 48 h, after which progesterone production was determined as described in Materials and Methods. Data are means of triplicate +/- standard error. These experiments were repeated four times.

### MEK inhibition stimulates expression of StAR

One of the processes that is involved in steroidogenesis is cholesterol transport from the outer to the inner mitochondrial membrane which is important for the conversion cholesterol into pregnenolone. Cytochrome P450scc participates in this processes as a rate-limiting enzyme and the whole process is regulated the steroidogenic acute regulatory (StAR) protein (7). The induction of StAR and its downstream effects are cAMP-dependent, as reported for gonadotropin-induced steroidogenesis in the gonads and ACTH-stimulated steroidogenesis in the fasciculata cells of the adrenal (7). Since StAR protein has a short functional half-life (28), we studied whether the ERK cascade is involved in the downregulation of StAR. For this purpose, rLHR-4 cells were treated with the various agents and examined for the expression of StAR 24 h after stimulation. As expected, LH, hCG, forskolin and to a considerably lesser extent dghCG induced StAR expression (but not ERK activation) under the conditions examined (Fig. 4). The MEK inhibitor PD98059 by itself caused an induction of StAR, but when this inhibitor was added with prior to forskolin, LH and hCG, there was a synergistic elevation in StAR expression. Similar results were obtained also in rFSHR-17 cells where MEK inhibition dramatically increased the forskolin- and FSH-induced expression of StAR. These results indicate that the ERK cascade negatively regulate steroidogenesis, as judged by the attenuation of StAR expression, which is the regulatory component that integrates the signals from both cAMP and ERK to regulate the rate of steroidogenesis.

**Fig. 4.**
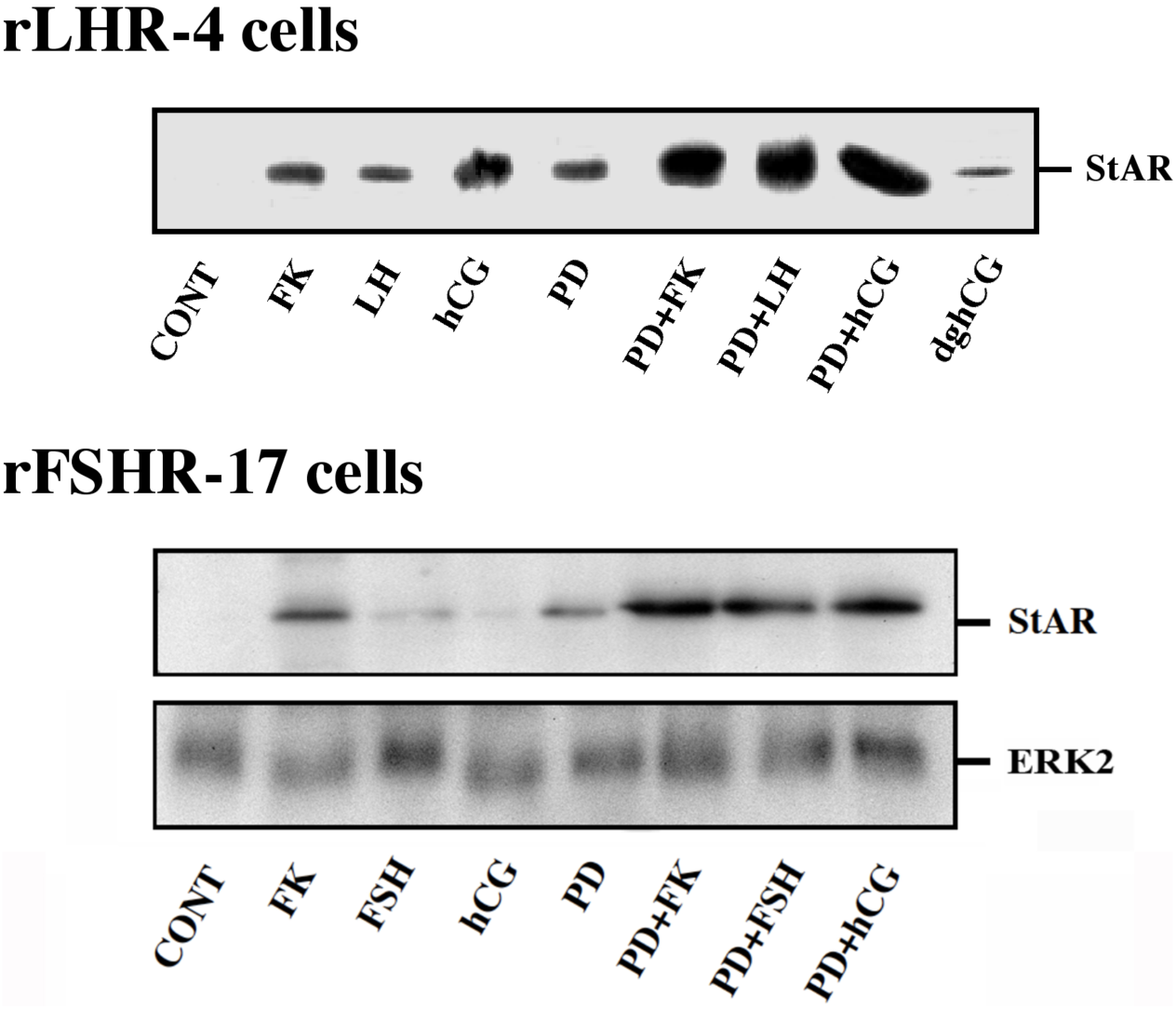
Expression of StAR in rLHR-4 and rFSHR-17 cells. Subconfluent cultures were stimulated with forskolin (FK, 50 *µ*M), PD98059 (PD, 25 *µ*M), hCG (3 iu/ml) hFSH (3iu/ml), dghCG (3 iu/ml), hLH (3 iu/ml) or combination of them as indicated for 24 h. Then, the cells were extracted as described under Materials and Methods and the extracts were subjected to SDS-PAGE and Western blotting using anti-StAR and anti-ERK antibodies. The arrow indicates mature StAR protein at 34 kDa. These experiments were repeated three times

PD98059 might have some unspecific effects that may influence the above results. To further verify them, we used another MEK inhibitor, the U0126 (29). Indeed, addition of this inhibitor to both rLHR-4 and rFSHR-17 cells caused an elevation in the amount of 30 kDa mature StAR (30) within 24 h (Fig. 5). The elevation was shown also by treatment of gonadotropins alone, but when the inhibitor was added together with a gonadotropin, the expression increased, reaching up to 10-fold above basal expression levels as well as U0126 and gonadotropin alone. We then studied the effect of U0126 on steroidogenesis in the rLHR-4 and the rFSHR-17 cells. Similarly to the results of PD98059, U0126 did not induce steroidogenesis by itself but synergized with the gonadotropins to produce high amounts of progesterone (Fig 5). Interestingly, in some of the experiments, a 37 kDa pre-StAR was detected (Fig. 5A). Usually, this cytosolic protein is maintained in a low level, because it rapidly matures into the 30 kDa form of StAR in the mitochondria (30). Unlike the 30 kDa StAR the amount of this protein remained low upon LH, FSH or MEK inhibition (Fig. 5).

**Fig. 5.**
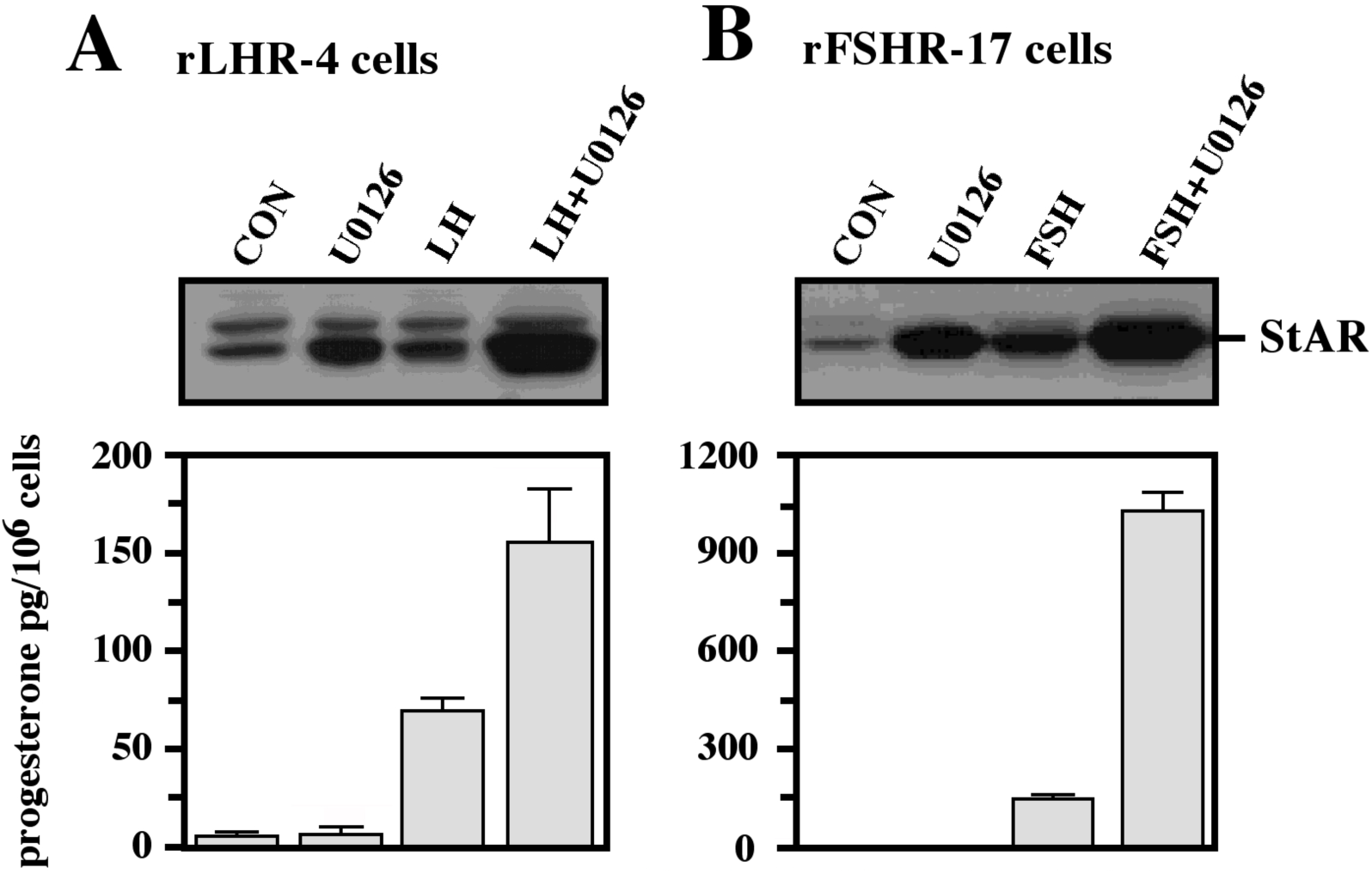
Effect of U0126 on StAR expression and progesterone production. Subconfluent cultures of either rLHR-4 (A) or rFSHR-17 (B) cells were stimulated for 24 h with LH (3 iu/ml, A) hFSH (3 iu/ml, B), U0126 (10 *µ*M) or combination of the gonadotropins with U0126 in the same concentrations. Expression of StAR (upper panel) and progesterone production (lower panel) were detected as described above.

Our results indicate that MEK inhibitors dramatically increase gonadotropin-induced StAR expression and steroidogenesis. They operate probably via elevation of StAR expression. The lack of corresponding elevation in progesterone production is probably due to the fact that in the immortalized granulosa cell lines there is no basal levels of the cytochrome p450scc that is obligatory for the conversion of cholesterol to pregnenolone (31). This notion is supported by our findings that in primary rat granulosa cells that do contain p450scc, PD98059 by itself increased progesterone production. On the other hand, MEK inhibitors do synergies with gonadotropin/cAMP stimulation of stereoidogenesis because of the de-novo synthesis of the cytochrome p450scc, which is stimulated by gonadotropin/cAMP in the granulosa cell lines (17,31)

### Subcellular localization of the overexpressed StAR

We then undertook to examine whether the elevated StAR by MEK inhibitors, gonadotropins and cAMP-elevating agents is mainly located in mitochondria (32). For this purpose, we stained rFSHR-17 cells with anti-StAR antibodies prior or following PD98059, FSH and forskolin stimulation (Fig. 6). As expected, no StAR expression was detected in non-stimulated cells. In contrast, StAR was evident in the mitochondria 24 h after treatment with PD98059. The same was seen after LH treatment, and this expression was dramatically increased in the mitochondria upon MEK inhibition. The same was seen for 8 Br cAMP alone, and its combination with PD98059. Thus, the immunocytochemical observations confirmed the data obtained by Western blot on the elevation of StAR expression, and showed that this elevation confined to the mitochondria.

**Fig. 6.**
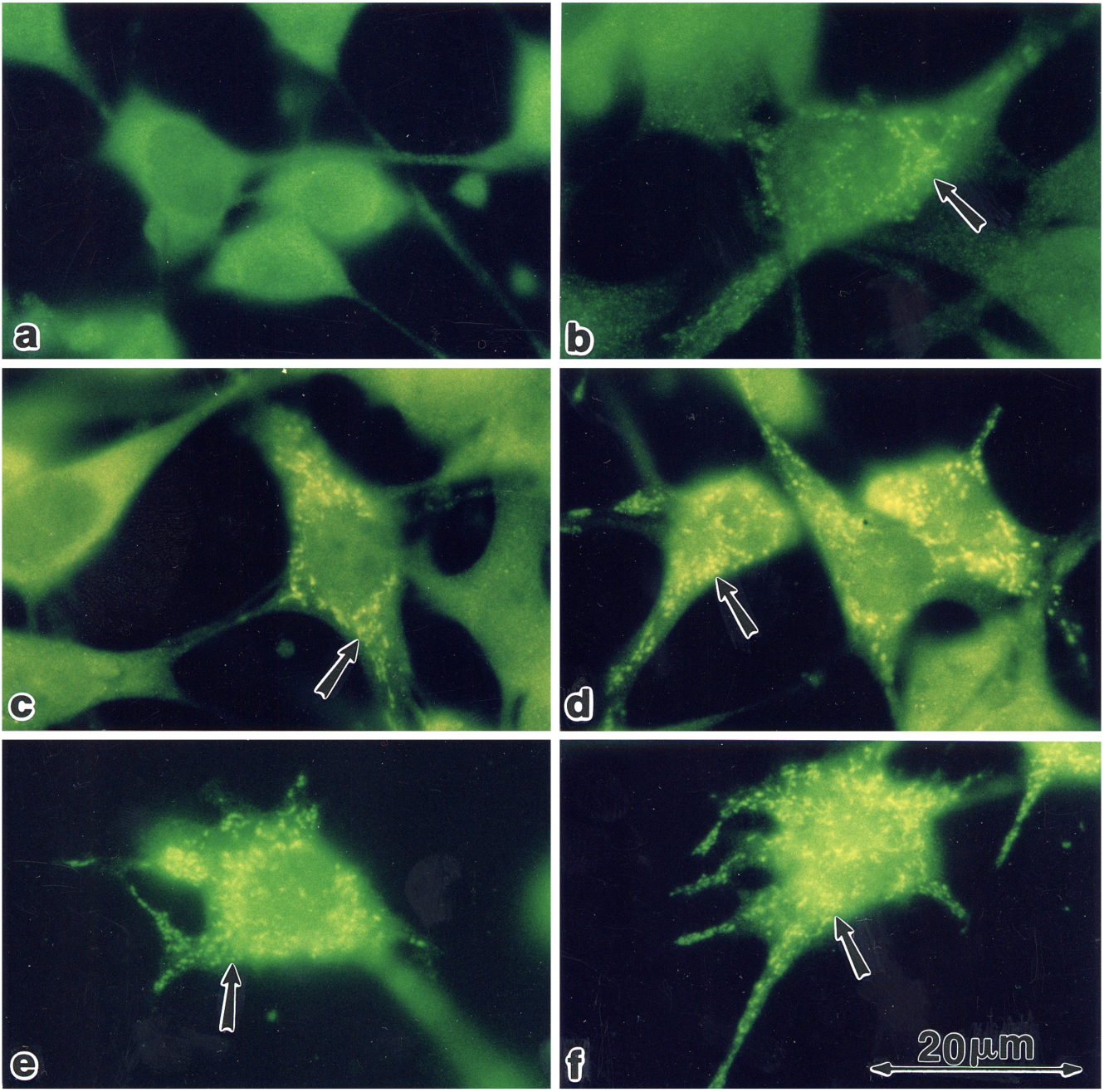
Subcellular localization of StAR upon induction with FSH and PD98059. Immunofloresence of cells stained with anti-StAR antibodies followed by goat anti-rabbit IgG conjugated to florescein, Subconfluent rFSHR-17 cells were stained with anti-StAR antibodies prior to or following PD98059, FSH and forskolin stimulation. a - no treatment; b - 24 h incubation with PD98059 (25 *µ*M); c - 24 h incubation with LH (3 iu/ml); d - 24 h incubation with PD98059 (25 *µ*M) and LH (3 iu/ml); e - 24 h incubation with 8-br-cAMP (50 *µ*M; f - 24 hr incubation with PD98059 (25 *µ*M) and 8-br-cAMP (50 *µ*M). Florescence microscopy x 1330. The arrow indicates StAR staining in the mitochondria.

### Gonadotropin-induced ERK activation and StAR production are mediated by PKA

Although we showed that an elevation of cAMP is sufficient to activate ERK, it was not clear whether cAMP and PKA are the major mediators of the gonadotropin-generated signaling to ERK. Therefore, we used H89, which is a potent and selective inhibitor of PKA. This inhibitor was used to study the involvement of cAMP/PKA in the activation of ERK in both rLHR-4 and rFSHR-17 cells. We found that 3 *µ*M of H89 15 min prior to gonadotropin stimulation did not change the basal phosphorylation of the three ERKs, but abrogated the induction of ERK by gonadotrophins (Fig. 7). As expected, ERK activation by forskolin and 8-Br-cAMP in both cell lines was also inhibited by H89 (data not shown), indicating that ERK activation is mediated mainly by PKA and probably not via the cAMP-GRF (33) or other means.

**Fig. 7.**
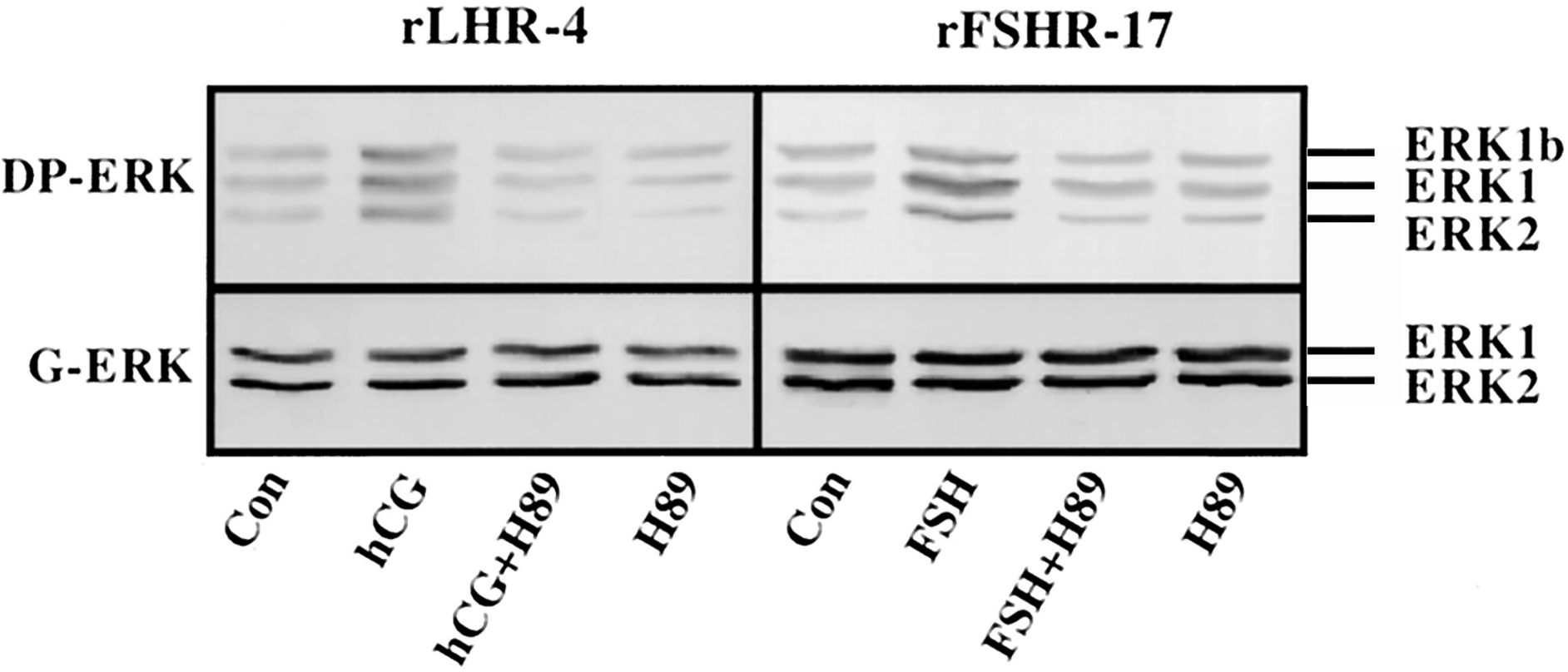
Effect of H89 on activation of ERK in gonadotropin-treated rLHR-4 and rFSHR-17 cells. rLHR-4 or rFSHr-17 cells were serum-starved for 16 h and then stimulated with the appropriate gonadotropins (3 iu/ml, 10 min) with or without the PKA inhibitor, H89 (15 min prestimulation, 3 *µ*M). Cytosolic extracts (50 *µ*g) were subjected to immunoblotting with DP-ERK (upper panel) or with anti-general ERK antibody (G-ERK, second panel). The ERK2, ERK1, and ERK1b are indicated.

In order to further verify the involvement of PKA in the activation of ERK by gonadotropins, we coexpressed GFP-ERK2 (21) together with the potent PKA inhibitor PKI or its inactive mutant (PKI^mutant^ (20)). The activation of ERK in the transfected cells was detected using the antibodies that recognize incorporation of phosphate into the activation loop of the GFP-ERK2. As observed with H89, PKI significantly inhibited ERK activation by gonadotropins and by forskolin (Fig. 8). These results clearly confirm that the activation of ERK by gonadotropin in the cell lines examined is mostly PKA dependent.

**Fig. 8.**
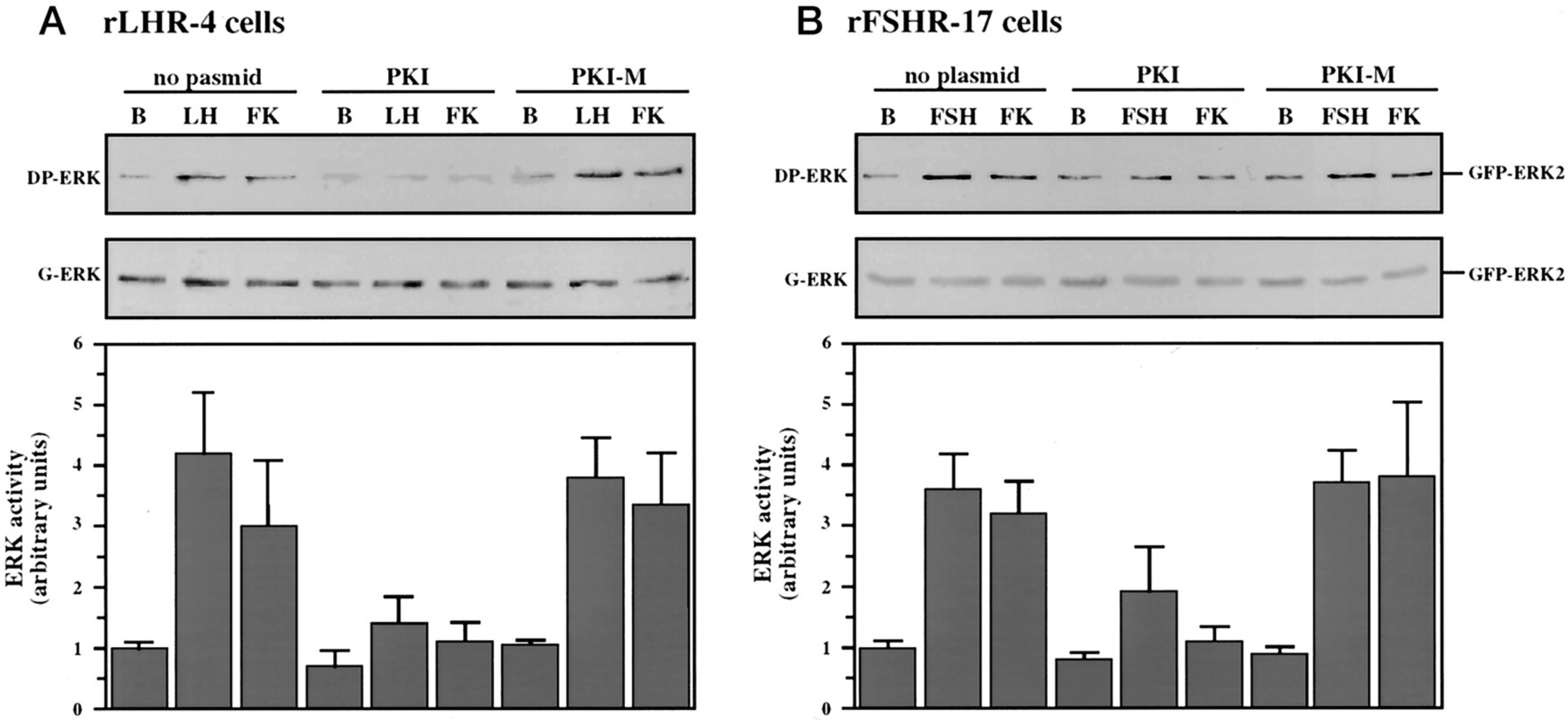
Effect of PKI on ERK-activation by gonadotropins and by forskolin. rLHR-4 (A) and rFSHr-17 (B) cells were transfected with pGFP-ERK2 alone (no plasmid) or cotransfected with pGFP-ERK2 together with RSV-PKI (PKI), and RSV-PKI^mutant^ ((PKI-M, which is inactive PKI). after transfection the cells were treated as described under Material and Methods for 18 h and then stimulated with FSH (3 iu), forskolin (FK, 50 *µ*M) for 10 min or left untreated (B). the cells were then harvested and cytosolic extracts were subjected to western blot analysis with the anti-DP-ERK and anti-C16 antibodies (G-ERK) the 70 kDa band which represent GFP-ERK2 is shown in the upper panels. Densitometric scanning of the DP-ERK lanes (arbitrary units) were used as a measure for ERK activity (bar graphs, bottom panels). The results in the bar graphs are average and standard errors of three experiments.

We then examined whether StAR activation is mediated by PKA alone or due other signaling pathways that cooperate with it. Indeed, H89 significantly inhibited hCG-FSH- and forskolin-stimulated StAR expression in rLHR-4 and rFSHR-17 cells (Fig. 9). These results indeed indicate that StAR production is regulated by PKA in the cell lines examined. As expected, progesterone production was also significantly inhibited by the H89 inhibitor (data not shown), indicating that the processes examined may function mainly downstream of PKA. ERK, although activated by PKA, serves as a negative regulator of this pathway due to its suppression of StAR (Fig. 10).

**Fig. 9.**
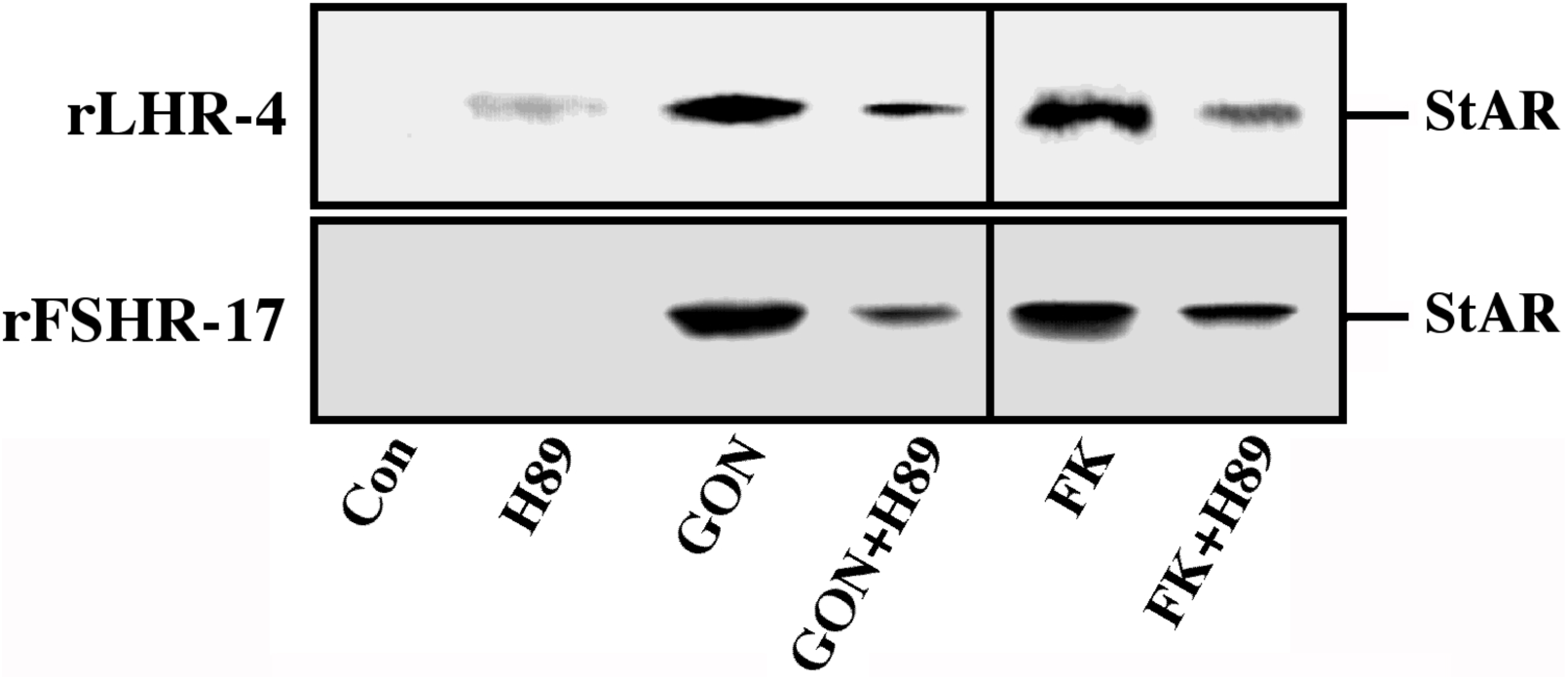
Effect of H89 on the expression of StAR in gonadotropin-treated rLHR-4 and rFSHR-17 cells. rLHR-4 or rFSHr-17 cells were serum-starved for 16 h and then stimulated with the appropriate gonadotropins (3 iu/ml, 10 min) with or without the PKA inhibitor, H89 (15 min prestimulation, 3 *µ*M). Cytosolic extracts (50 *µ*g) were subjected to immunoblotting with anti-StAR antibody. The arrow indicates mature StAR protein at 32 kDa.

**Fig. 10.**
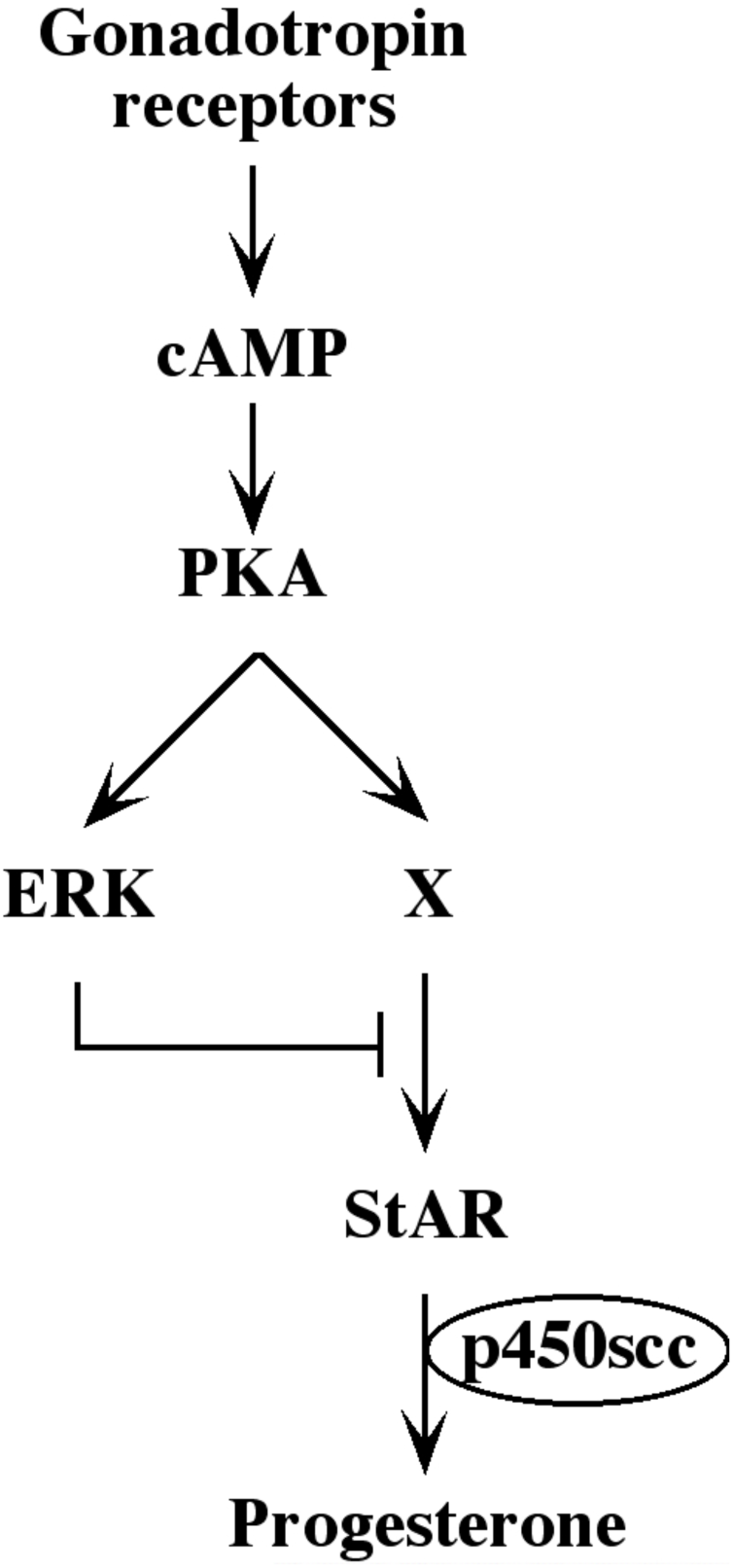
Schematic representation of the signaling pathways controlling gonadotropin - induced steroidogenesis.

## Discussion

Crosstalk between signaling pathways is an important mechanism to fine-tune the stimulated outcomes. In this study we demonstrate a mechanism of cross-talk between the cAMP/PKA and the ERK cascade in a Gs-induced system. The interaction between these two cascades has already been studied in several cellular systems, revealing different interactions and effects (34). For example, it was shown that in EGF-stimulated Rat1 fibroblasts (35) or PDGF-stimulated human arterial smooth muscle cells (36) cAMP inhibits the activation of the ERK cascade. This inhibition was proposed to occur by either inhibitory phosphorylation of Raf-1 (35) or by activation of the small GTPase, Rap-1, which compete with Ras for the activation of Raf-1 (37). On the contrary, in NGF-stimulated PC12 cells, cAMP not only does not inhibit ERK, but in fact activates it, in order to induce mitogenic or differentiation processes. Several mechanisms have been proposed for the activation of ERK downstream of cAMP. One of them is the activation of the cAMP responsive guanine-nucleotide exchange factors for the small GTPase Rap1, Epac1 and Epac2 (33). Thus, upon binding of cAMP, these components activate Rap1, which then promotes the activation of B-Raf, leading to the activation of the rest of the ERK cascade. However, in the rLHR-4 and rFSHR-17 cells studied here, the activation of ERK sis downstream of PKA, indicating that the Epac’s are probably not involved in it. The activation downstream of PKA may involve an activation of the Rap-1 GTPase, which in turn associate with B-Raf. Interestingly, another possibility for such a crosstalk is the activation of the cAMP-responsive STE-20-like kinase, MST3b, that was shown to cause activation of ERK by cAMP in the brain (38). However, this specific isoform does not seem to be expressed in granulosa cells and its connection to PKA is not known. However, it is possible that another MAP4K or MAP3K is involved in the transmission of PKA signals to ERK in the rLHR-4 and rFSHR-17 cells.

In this study, we demonstrated the involvement of PKA in gonadotropin-dependent ERK activation both by pharmacological means using PKA inhibitor H89, and by transfection of cells with plasmid encoding for PKI. The results obtained by both methods indicate that PKA plays a major role in transducing the gonadotropin signals towards ERK. Nevertheless, it should be noted that although PKI completely suppressed forskolin-induced ERK activation, it did not completely inhibit the gonadotropin-induced ERK activation. Therefore it is possible that the LH and FSH receptors use other G proteins or the Gβγ to activate the ERK cascade as was observed for other receptors and cell types (reviewed in (23,39). Interestingly we recently found that bFGF suppresses progesterone production in the granulosa cell lines (data not shown). This suggests that there is an alternative pathways in these cells that suppress steroidogenesis via ERK, which is not stimulated by the cAMP/PKA pathway.

As mentioned above, cooperation between the cAMP/PKA pathway and the ERK cascade has been demonstrated in several cells and systems. For example, it was shown that cAMP causes sustained activation of the ERK cascade, important for neurite outgrowth in PC-12 cells (40). In human cyst epithelial cells, cAMP causes a mitogenic response via the ERK cascade (41). Interestingly, cAMP might contribute to a late down-regulation of ERK-mediated processes. An example for such interaction is the PKA-induced CPG16 kinase, which seems to partially inhibit the activity of the transcription factor CREB (42). This suggests involvement of the down-regulation of cAMP- and ERK cascade-induced transcription. In contrast, here we show that the activation of processes downstream of PKA may also be inhibited by an ERK-mediated mechanism.

The inhibition progesterone production downstream of cAMP might be regulated by phosphorylation-dephosphorylation of proteins that play a role in the steroidogenesis. Here we examined the StAR protein that is known to be phosphorylated on serine or threonine residues (43). However, the expression of StAR did not correlate with the induction of PKA or ERK cascades. In addition, we did not detect any direct phosphorylation of StAR by ERK (data not shown). On the other hand, we did observe an inverse correlation between ERK activity and StAR expression. Indeed, blockade of ERK activity caused an elevated StAR protein expression, while activation of the kinase reduced StAR expression in granulosa cells. Therefore, it seems that the two cascades interact to regulate StAR transcription, which is the primary mechanism that regulate StAR expression in granulosa cells (44). It was previously reported that the transcription of StAR is induced by several transcription factors, including steroidogenic factor-1 (SF-1), C/EBP and the negative regulator DAX-1 (45-47). SF-1 and C/EBP are probably regulated downstream of PKA. However, these components are unlikely to participate in the down-regulation of StAR expression via the ERK cascade. This is because both have been shown to be stimulated and not inhibited by ERK (48,49). Therefore, it is possible that the negative regulation of StAR expression occurs by DAX-1 or by another, un-identified transcription factor. Alternatively, the reduction of StAR expression could be controlled by induction of potent phosphatases that abolish both the PKA and ERK phosphorylation of SF-1 and C/EBP. In addition, it could be regulated by the proteolytic system that reduces the half-life of the StAR protein. Another explanation is the involvement of the signaling pathway in desensitization of the gonadotropin receptors. Indeed, prolonged exposure of granulosa cells to gonadotropic hormones was shown to cause desensitization of the cells to further stimulation. This is characterized by downregulation of cAMP formation as well as of steroidogenesis (9). Moreover, it was also demonstrated that ERK may activate G-protein coupled receptor kinase 2 (50), which in turn induces down-regulation of the seven transmembrane receptors (GPCRs). However, we don’t think that this is the mechanism here, because the inhibitory effects of ERK were also demonstrated when cells were stimulated by cAMP-inducing agents. Since this activation of cAMP bypasses the receptor in activating PKA, we believe that most of the inhibitory signals are probably receptor-independent. Nevertheless, it is very likely that under physiological conditions the gonadotropins are the key stimulators of ERK activity. Finally, ERK activation can explain the mitogenic signals that were detected upon FSH stimulationduring folliculogenesis.

The initiation of steroidogenesis, is proportional to the duration and extent of cAMP production. However, we found that full activation of ERK can be achieved as a consequence of even modest increases in intracellular cAMP. This amplification process probably occurs by a switch-like mechanism of the ERK cascade, which allows a strong signaling output even by weak extracellular signals (51). The high activity of ERK, which functions downstream of cAMP, may explain the suppression of steroidogenesis upon weak gonadotropic signals, which lead to steroidogenesis. Therefore, it is likely that this situation causes the low levels of steroidogenesis induced by dghCG, which is able to induce only weak signals by the gonadotropic receptors.

In summary, we show here that activation of cAMP/PKA signaling by gonadotropins not only induces steroidogenesis, but also activates down-regulation machinery, which in our case involves the ERK cascade. This negative regulation inhibits the gonadotropin-induced steroidogenic pathway. The mechanisms that are involved in the inhibition are different from the well-characterized receptor desensitization. ERK activation downstream of PKA, might in turn regulate StAR expression, which is a key factor in the down-regulation processes. Thus, cAMP/PKA not only induces gonadotropic-induced steroidogenesis, but it also activates the downregulation mechanism to silence steroidogenesis under certain conditions. Our findings also raise the possibility that modulation of the activity of the ERK cascde by other pathways is an important mechanism for diminution or amplification of gonadotropin-stimulated steroidogenesis. In this sense, this inhibition may regulate functional luteolysis, which is a process in which leutinized granulosa cells show reduced sensitivity to LH despite maintenance of LH receptor or to up-regulation of the steroidogenic machinery during lutinization of granulosa cells (reviewed in (52)).

## Acknowledgments

This work was supported by grants from the Benozyio Institute for Molecular Medicine at the Weizmann Institute of Science and from the Estate of Siegmund Landau to RS and the Israel Academy of Science to AA, and NIH grant HD-06224 to JFS. AA is the incumbent of the Joyce and Ben. B. Eisenberg professorial chair in molecular endocrinology and cancer research.

\# - This is a correction of the withdrawn paper: J Biol Chem. 2001 276(17):13957-64. Withdrawal notice: J Biol Chem. 2017, 292(21):8847. The problematic figure 4 was corrected.

## Abbreviations

DP-ERK Ab: anti-diphospho-ERK antibody;
dghCG: deglycosilated hCG;
ERK: extracellular signal-regulated kinase;
FSH: follicle stimulating hormone;
hCG: human chorionic gonadotropin;
MAPK: mitogen-activated protein kinase;
MBP: myelin basic protein;
LH: luteinizing hormone;
PKA: protein kinase A;
SF-1: steroidogenic factor-1;
StAR: steroidogenic acute regulatory protein;
TPA: tetradecanoyl phorbol acetate.

